# Conformal Uncertainty Quantification for BayesAge Epigenetic Age Predictions

**DOI:** 10.64898/2026.08.11.742144

**Authors:** Megan Mitchell, Lajoyce Mboning, Louis-S. Bouchard, Matteo Pellegrini

## Abstract

Epigenetic clocks predict chronological age from DNA methylation (DNAm) profiles, yet most provide point estimates without calibrated uncertainty. This limits their use when quantified error bounds are required. We apply conformal prediction to BayesAge, a maximum-likelihood clock that models nonlinear DNAm–age relationships using a small set of CpG loci and a count-based likelihood. Split conformal prediction yields distribution-free prediction intervals with finite-sample marginal coverage guarantees under exchangeability and requires only a single model fit per split. We also evaluate a locally scaled variant that produces age-dependent interval widths using a locally estimated error scale. In our targeted bisulfite sequencing cohort, split conformalized BayesAge attains near-nominal empirical coverage while preserving BayesAge point-prediction accuracy. The locally scaled variant yields wider intervals at older ages, but its coverage is less stable in small-calibration regimes, consistent with additional uncertainty from estimating the local scale. Relative to Monte Carlo intervals that propagate read-sampling variability and to higher-dimensional linear baselines (conformalized linear quantile regression and conformalized Lasso regression), conformalized BayesAge provides calibrated uncertainty using substantially fewer CpG sites and with weaker age-dependent structure in residuals. These results support conformal prediction as a practical approach for uncertainty quantification in DNAm-based age estimation.

---

Although an individual’s DNA sequence is largely stable throughout life, the epigenome changes with age and environment. One widely studied epigenetic mark is DNAm, the covalent addition of a methyl group to the 5’ carbon of cytosine, most commonly at cytosine-phosphate-guanine (CpG) dinucleotides, forming 5-methylcytosine. DNAm contributes to transcriptional regulation and is required for normal development (1). DNAm levels also vary across individuals because of environmental exposures, lifestyle, cell-type composition, and stochastic processes, a phenomenon often termed epigenetic drift (2, 3). DNAm levels at specific CpG sites correlate with chronological age and have been used to construct epigenetic clocks that predict chronological age from DNAm profiles (2, 4–6). The difference between a clock-predicted age and chronological age is commonly referred to as age acceleration. Age acceleration has been associated with mortality and multiple age-related diseases in many cohorts, although associations depend on study design and confounder control (2–4, 7).

Early widely used clocks developed by Horvath (5) and Hannum (2) employed penalized multivariate linear regression on panels of CpG sites (353 and 71 sites, respectively). In healthy cohorts, these models often achieve mean absolute error (MAE) values on the order of a few years, with performance generally improving as training set size increases (5, 8). However, high-dimensional linear models can be sensitive to cohort shift and technical variation, including batch effects, and can show degraded precision or calibration when applied to independent datasets (9). Discrepancies between clocks have also been reported in perturbed settings such as cellular reprogramming, where measurement and batch effects can be substantial (10). In addition, most clocks report point estimates without calibrated uncertainty (2, 5, 6). This limits their interpretability in applications that require quantified error bounds. Finally, linear predictors do not explicitly model nonlinear DNAm– age trajectories, despite evidence that many CpG sites exhibit nonlinear patterns across the lifespan (11–13).

We previously introduced BayesAge, a maximum-likelihood approach that predicts chronological age from a small subset of CpG loci and accommodates nonlinear DNAm–age relationships through locally weighted scatterplot smoothing (LOWESS) together with a binomial count likelihood (14). In that work, uncertainty was estimated by Monte Carlo simulation of DNAm counts at each locus followed by propagation through the BayesAge predictor. This procedure is computationally intensive and, in our experience on aggregated datasets, did not yield calibrated empirical coverage, consistent with the fact that it primarily propagates read-sampling variability and does not directly account for model error or cohort heterogeneity.

Conformal prediction provides an alternative. It wraps around any point predictor and yields finite-sample marginal coverage guarantees under exchangeability, without requiring a parametric error model (15). Conformal methods have been applied to epigenetic clocks, including conformalized linear quantile regression in childhood datasets (16). Such approaches can produce interval widths that adapt to heteroskedastic prediction errors, which may arise from age-dependent variability in DNAm (13). However, conformalized linear quantile regression requires a separate quantile model and is not directly aligned with the BayesAge likelihood-based estimator.

Here we study two conformal approaches that apply directly to BayesAge: split conformal prediction and a locally scaled variant based on locally estimated error scales. Split conformal prediction is computationally efficient, can be used with any underlying age predictor, and provides finite-sample marginal coverage guarantees under exchangeability (15). The locally scaled variant uses a rescaled nonconformity score to adapt interval widths to local variability and to reflect heteroskedasticity, but its finite-sample coverage can be less stable when calibration sets are small (17, 18). We find that split conformal prediction improves empirical coverage relative to simulation-based intervals while preserving BayesAge point-prediction accuracy. The locally scaled method produces age-dependent interval widths that track variation in absolute error across age, but exhibits weaker coverage control in small-calibration regimes.

## Results

### Split conformal prediction and empirical coverage

Split conformal prediction allocates a subset of labeled samples to a calibration set in order to estimate a residual quantile used to form prediction intervals. This allocation can reduce the effective training size for the underlying point predictor and, in principle, can increase point-prediction error. We therefore first evaluated BayesAge point prediction without conformalization. For this point-estimation baseline, we used the same validation partitioning described in Materials and Methods, but fit BayesAge on all non-validation samples in each split (i.e., without reserving a calibration subset). Using 16 CpG sites, BayesAge attained a mean absolute error (MAE) of 6.098 years.

We next applied split conformal prediction with the same CpG-selection procedure and the same validation partitioning, using *m* = 200 calibration samples per split and target marginal coverage *δ* = 0.9. Relative to the point-prediction-only analysis, this reduced the number of samples used to fit the BayesAge reference curves by *m* per split (from 900 to 700 samples in the first 11 splits, and from 968 to 768 samples in the final split). Despite this reduction, conformalized BayesAge attained a similar MAE of 6.083 years on held-out validation samples.

We define empirical coverage as the fraction of validation samples for which the true chronological age lies within the reported prediction interval. For *δ* = 0.9, split conformalized BayesAge achieved an empirical coverage of 91.2%, close to the nominal level. Before bounding to the admissible age range, the average half-width was 12.984 years (intervals of the form [*ŷ* − Δ, *ŷ* + Δ] with Δ = 12.984 on average across splits). For reporting, we bounded each interval to [0, 100] years, which can induce asymmetry near the boundaries; after bounding, the mean total width was 25.61 years.

To probe behavior in a small-calibration regime, we also performed a stress test using 10 training samples, 10 calibration samples, and 10 validation samples over 1000 random splits. In this setting, the running average empirical coverage stabilized near the target level, consistent with split conformal prediction’s finite-sample marginal coverage guarantee under exchangeability (Fig. 3).

**Fig. 1.**
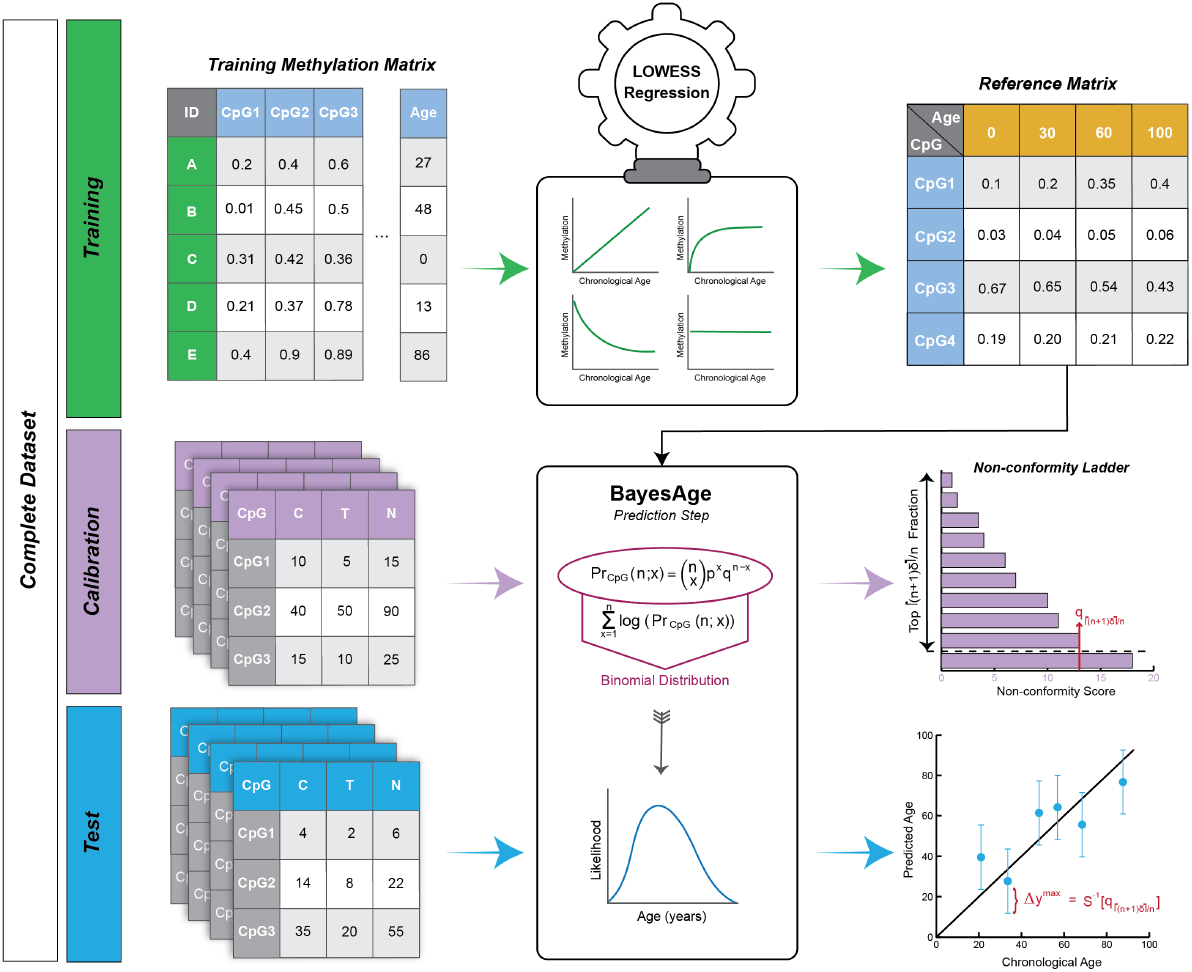
Workflow for BayesAge with split conformal prediction. The full dataset contains 988 samples. We performed 12 splits. In each split, 200 samples were used for calibration and the remaining samples were divided between training and validation. In the first 11 splits 700 samples were used for training and 88 were used for validation. In the final split only the remaining 20 samples were used for validation, and the remaining samples were used for training. Each sample was used as a validation point exactly once across the 12 splits. For each split, we fit BayesAge on the training set to construct the reference matrix, predicted ages for calibration samples, and computed nonconformity scores as absolute errors on the calibration set. We then formed split conformal prediction intervals for validation samples using the calibration quantile and bounded intervals to the admissible age range [0, 100].

**Fig. 2.**
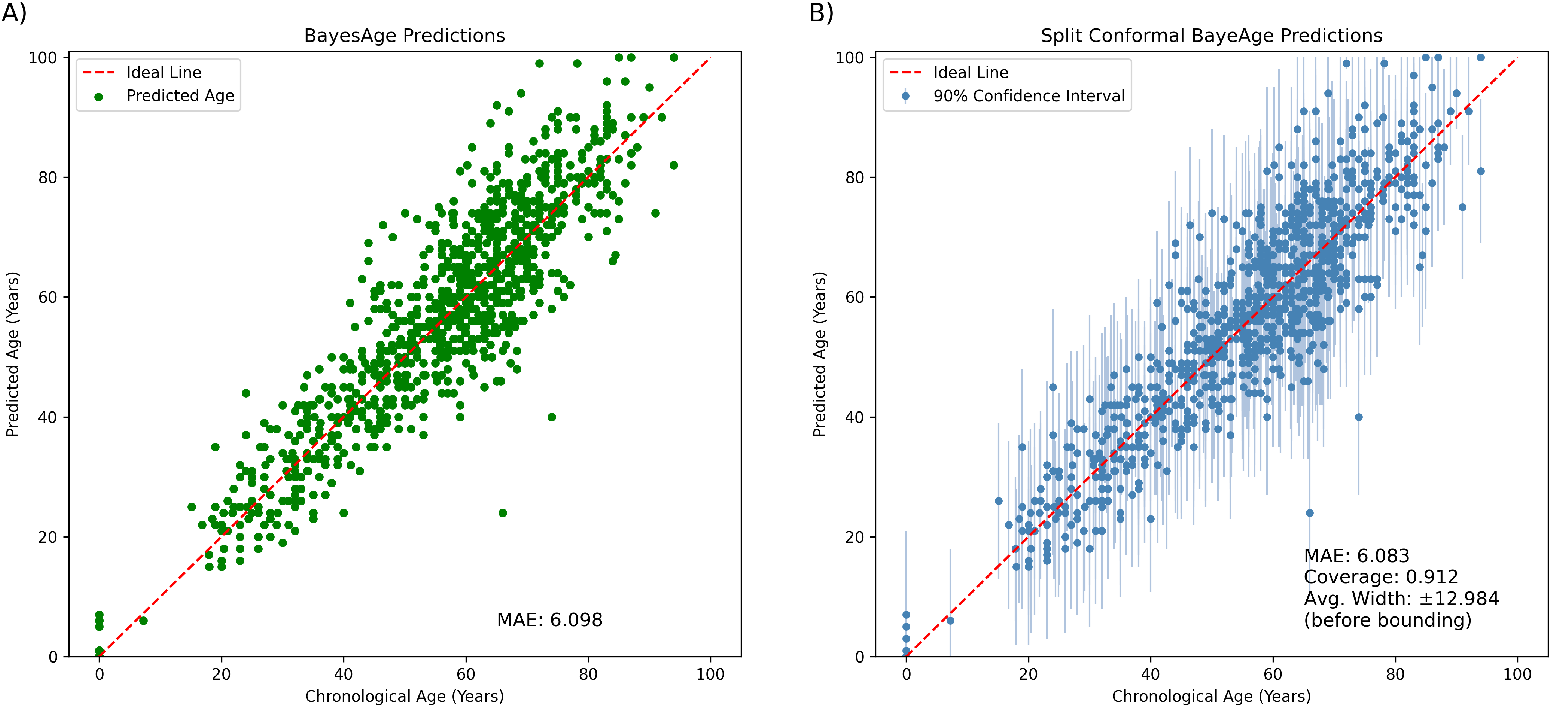
Chronological age predictions from DNAm using BayesAge with 16 CpG sites. **A)** BayesAge point predictions under the cross-validation protocol described in Materials and Methods (validation sets of 88 samples per split, besides 20 used for the final split; remaining samples used for training). **B)** BayesAge with split conformal prediction intervals using 200 calibration samples per split. Points show predicted ages; intervals reflect bounded split conformal prediction intervals at target coverage *δ* = 0.9.

**Fig. 3.**
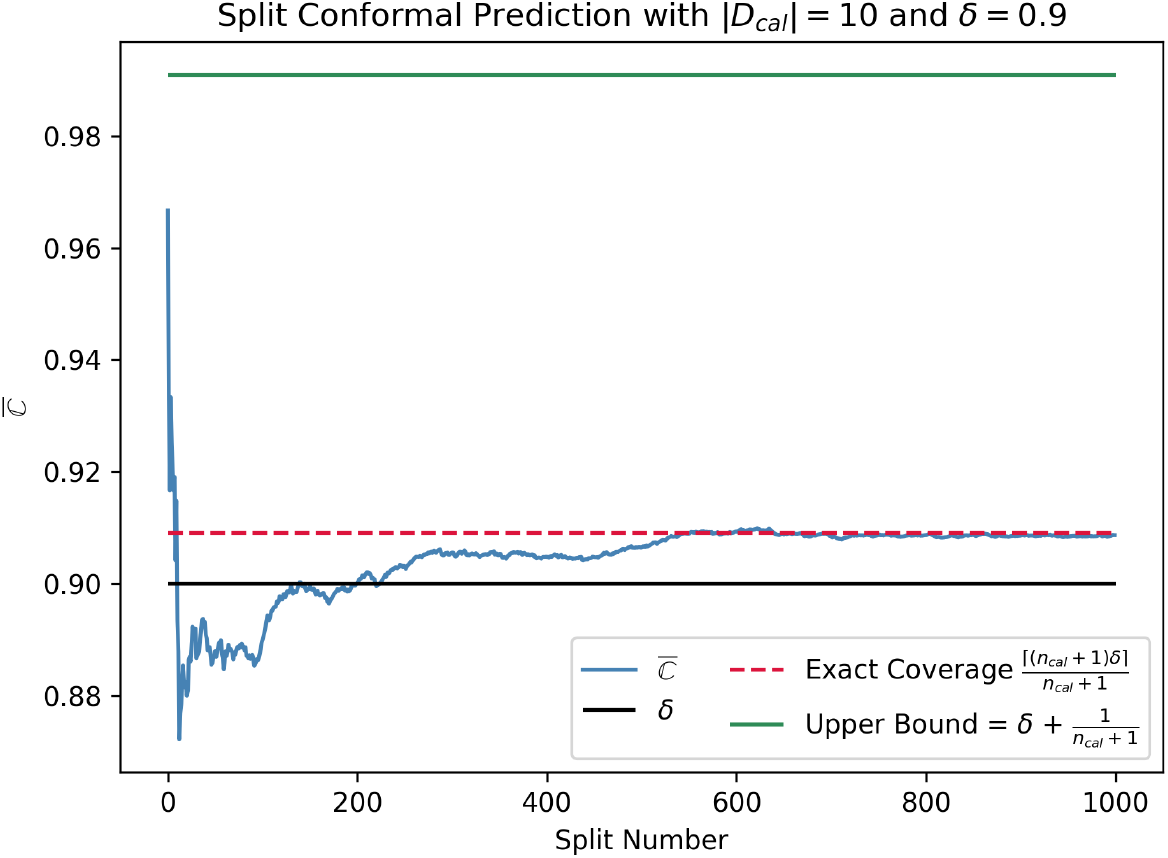
Small-sample stress test for split conformal prediction. We performed 1000 random splits of 30 samples into 10 training, 10 calibration, and 10 validation examples at target coverage *δ* = 0.9. The curve shows the running average of empirical coverage across splits, defined as the fraction of validation examples whose true ages fall within the reported prediction intervals.

**Fig. 4.**
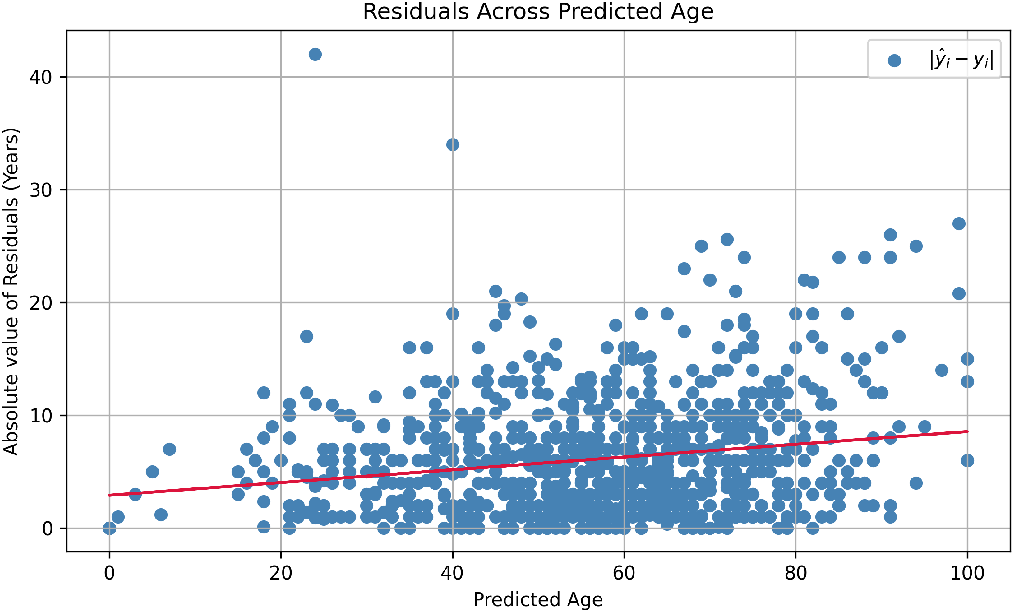
Absolute residuals |*ŷ* − *y*| versus predicted age *ŷ* for split conformalized BayesAge. The signed residual (age acceleration) is *ŷ* − *y*, and the plotted values are its absolute value. The smooth curve summarizes the dependence of absolute error on predicted age.

### Locally scaled conformal prediction and age-dependent interval widths

Split conformal prediction uses a single calibration quantile per split and therefore produces constant-width intervals within that split prior to bounding. DNAm-based age prediction error can plausibly be heteroskedastic across age, due to age-dependent variability in DNAm and differences in measurement or cell-type composition across age ranges (2). Motivated by this possibility and by the observed residual patterns, we evaluated a locally scaled conformal variant in which the nonconformity score divides the absolute residual by a locally estimated error scale, yielding prediction intervals whose widths vary with predicted age (17, 18).

Using the same cross-validation protocol, CpG selection, and *m* = 200 calibration samples per split, and again targeting *δ* = 0.9, the locally scaled method achieved an empirical coverage of 89.6% and an average bounded total width of 24.868 years. The resulting bounded intervals tended to widen with increasing predicted age (Fig. 5), consistent with age-dependent variation in absolute error.

**Fig. 5.**
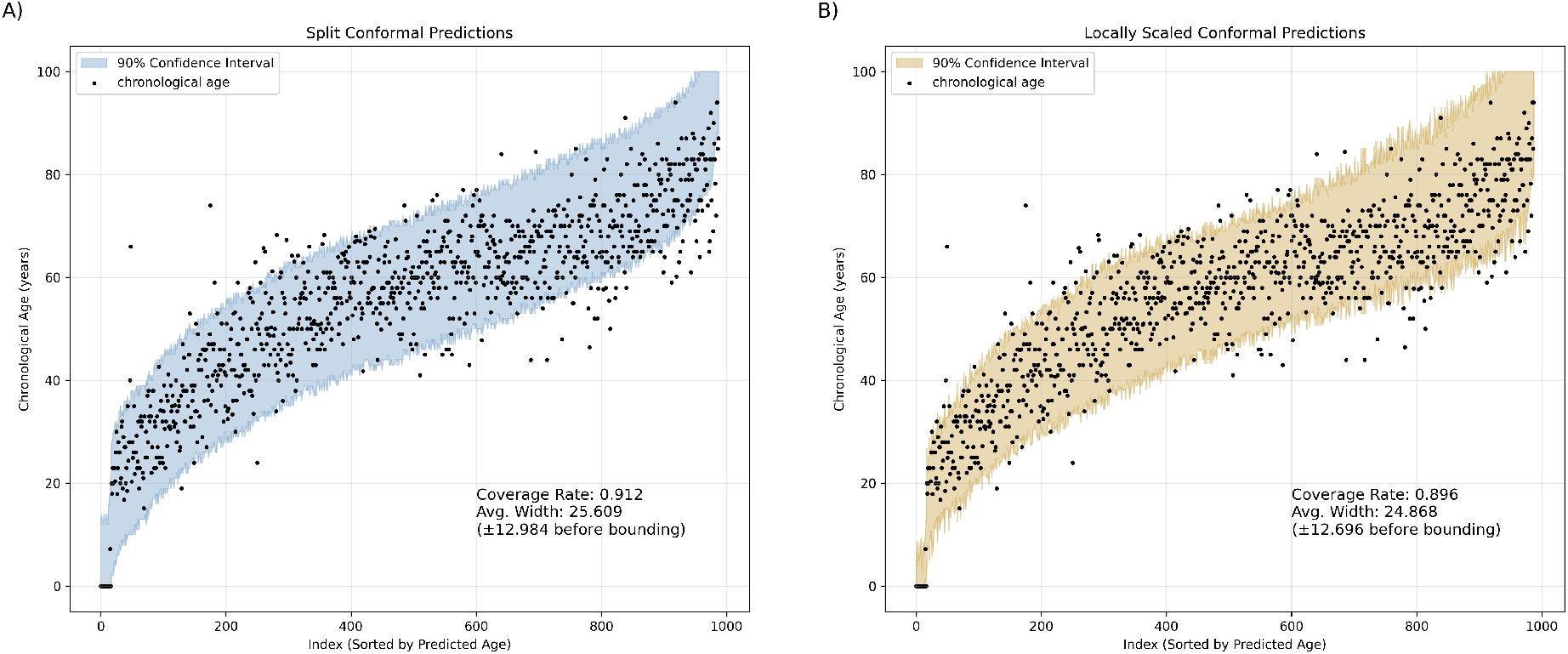
Prediction intervals for split conformal and locally scaled conformalized BayesAge at target coverage *δ* = 0.9 with 16 CpG sites and 200 calibration samples per split. **A)** Split conformal prediction intervals. **B)** Locally scaled conformal prediction intervals. Samples are sorted by predicted age. Shaded regions denote bounded prediction intervals [*L*_*i*_, *U*_*i*_] ⊆ [0, 100].

To quantify this pattern, we summarized interval widths within age bins of width 14 years (Fig. 6). Split conformal intervals were narrower for younger samples, an effect partly driven by bounding at age 0, and were otherwise relatively stable across midlife. In contrast, locally scaled intervals increased in width with age, reflecting the local scale estimate.

**Fig. 6.**
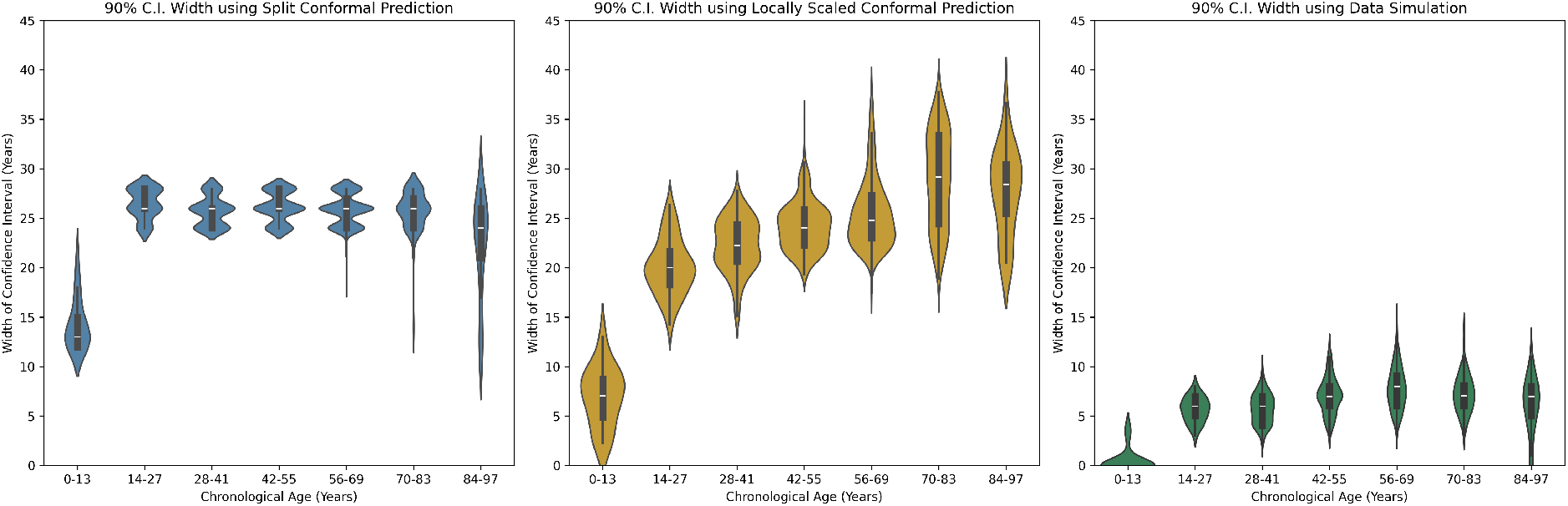
Interval widths as a function of age. Box plots summarize the distribution of bounded interval widths (*U*_*i*_ − *L*_*i*_) within 14-year bins, for split conformalized BayesAge, locally scaled conformalized BayesAge, and Monte Carlo simulation-based intervals.

Unlike split conformal prediction, the locally scaled method does not generally retain the same finite-sample marginal coverage guarantee unless the scale function is estimated independently of the calibration scores. In our implementation, the scale is estimated from calibration residuals, which introduces additional randomness and can weaken coverage control when calibration sets are small. This limitation is apparent in the small-sample stress test: with 10 training, 10 calibration, and 10 validation samples over 1000 random splits, the empirical coverage did not approach the nominal level (Fig. 7).

**Fig. 7.**
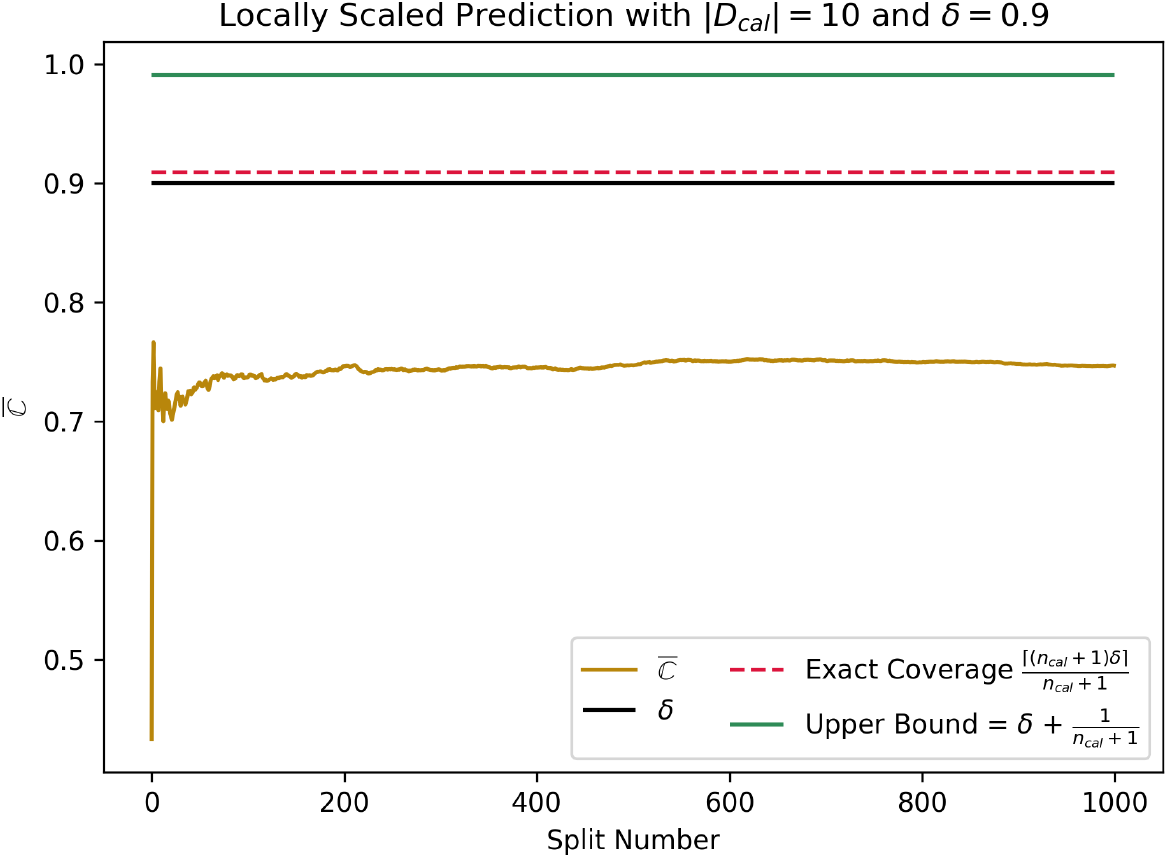
Small-sample stress test for locally scaled conformal prediction. We performed 1000 random splits of 30 samples into 10 training, 10 calibration, and 10 validation examples at target coverage *δ* = 0.9. The curve shows the running average of empirical coverage across splits, defined as the fraction of validation examples whose true ages fall within the reported prediction intervals.

Finally, we compared both approaches across multiple target coverage levels using *m* = 200 calibration samples and 16 CpG sites. Across *δ* ∈ {0.5, …, 0.99}, split conformal prediction achieved empirical coverage closer to nominal while producing interval widths comparable to those of the locally scaled method (Table 1).

**Table 1.**
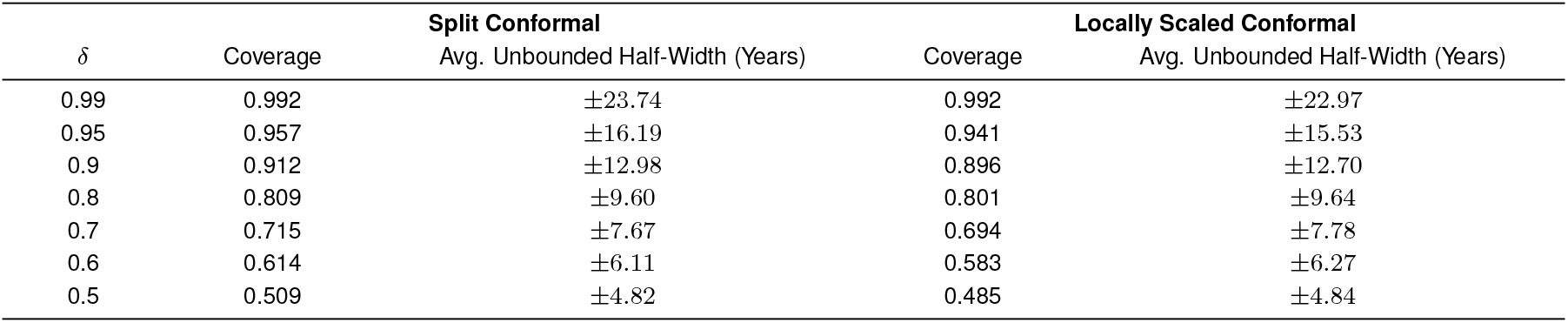
Prediction Interval Performance at Different Target Coverage Levels. Coverage is the empirical fraction of validation examples whose true ages fall within the reported prediction intervals, pooled across splits. “Avg. Unbounded Half-Width” denotes the mean interval half-width before truncation to [0, 100]: *q*\* for split conformal prediction, and 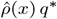 averaged over validation examples for locally scaled conformal prediction.

**Table 2.**
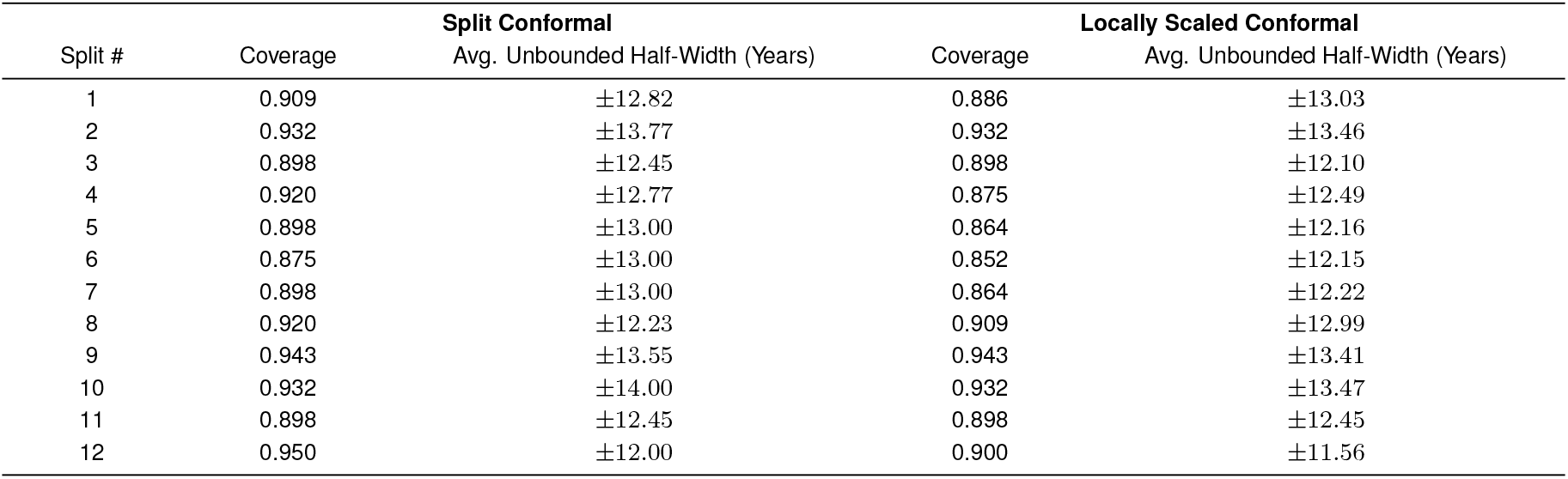
Prediction Interval Performance Across Splits with *δ* = 0.9.

### Comparison with alternative uncertainty quantification base-lines

We compared conformalized BayesAge with three baseline approaches to uncertainty quantification at target marginal coverage *δ* = 0.9: (i) Monte Carlo simulation-based intervals as in the original BayesAge implementation, (ii) conformalized linear quantile regression implemented using a linear `QuantileRegressor` with conformal calibration (hereafter, conformalized linear quantile regression), and (iii) split conformal prediction wrapped around Lasso regression. For each method, empirical coverage was computed as the fraction of validation samples whose true chronological age fell between the reported lower and upper bounds. Reported interval widths correspond to the mean of (*U*_*i*_ − *L*_*i*_) over validation samples, where [*L*_*i*_, *U*_*i*_] denotes the bounded interval after truncation to the admissible age range [0, 100]; bounding can induce asymmetry around the point prediction.

Both split conformal and locally scaled conformalized BayesAge achieved empirical coverage close to nominal, with mean widths comparable to those of the conformalized linear quantile regression baseline (Fig. 8). The conformalized linear quantile regression baseline achieved 90.3% empirical coverage with a mean bounded width of 23.18 years. In contrast, simulation-based intervals exhibited substantial undercoverage (42.3%), consistent with intervals that are too narrow when uncertainty sources beyond read-sampling variability are present. Split conformal Lasso regression produced the narrowest intervals among the evaluated methods (mean width 16.33 years) while maintaining near-nominal coverage (90.2%).

**Fig. 8.**
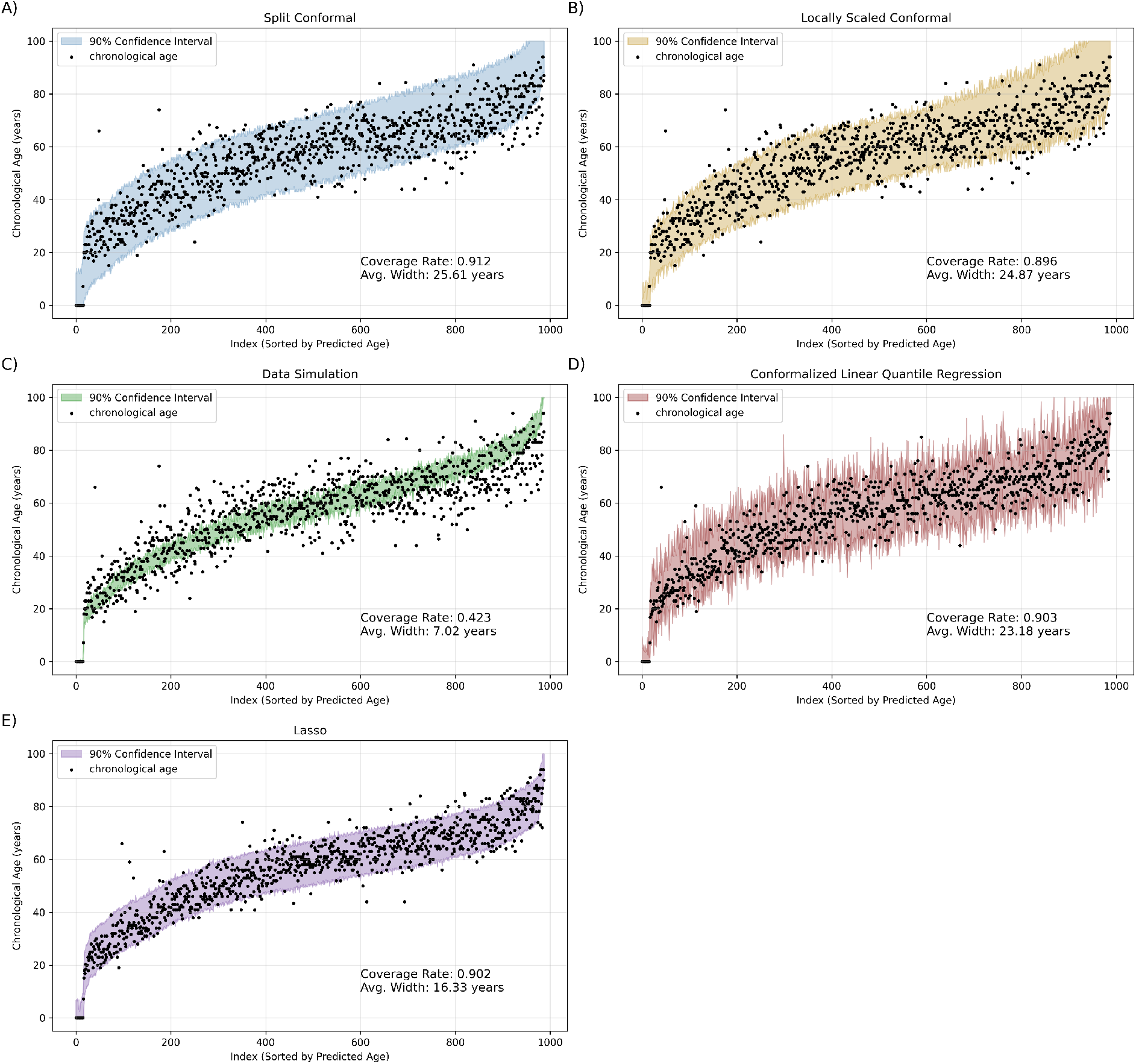
Prediction interval coverage across uncertainty quantification methods at target coverage *δ* = 0.9. Samples are sorted by predicted age and the shaded region denotes the bounded prediction interval [*L*_*i*_, *U*_*i*_] ⊆ [0, 100]. **A)** Split conformalized BayesAge. **B)** Locally scaled conformalized BayesAge. **C)** Monte Carlo simulation-based intervals using 100 synthetic draws per sample. **D)** Conformalized linear quantile regression implemented using `MapieQuantileRegressor` with a linear `QuantileRegressor`. **E)** Split conformalized Lasso regression with *λ* = 0.02 (the `scikit-learn` parameter `alpha`).

We also examined residual structure as a function of age, using signed age acceleration (predicted age minus chronological age) as the residual. In this dataset and implementation, BayesAge residuals show less age-dependent structure than the Lasso and conformalized linear quantile regression baselines, which display increasing negative residuals at older ages (Fig. 9). Because conformal prediction wraps a point predictor without changing its point predictions, these residual patterns reflect differences among the underlying predictors rather than an effect of conformalization. While conformalized BayesAge provides calibrated uncertainty with substantially fewer CpG sites, it yields higher point-prediction error than the higher-dimensional linear baselines in this study: the conformalized BayesAge MAE was 6.083 years, compared with 4.694 years for conformalized linear quantile regression and 3.925 years for conformalized Lasso regression.

**Fig. 9.**
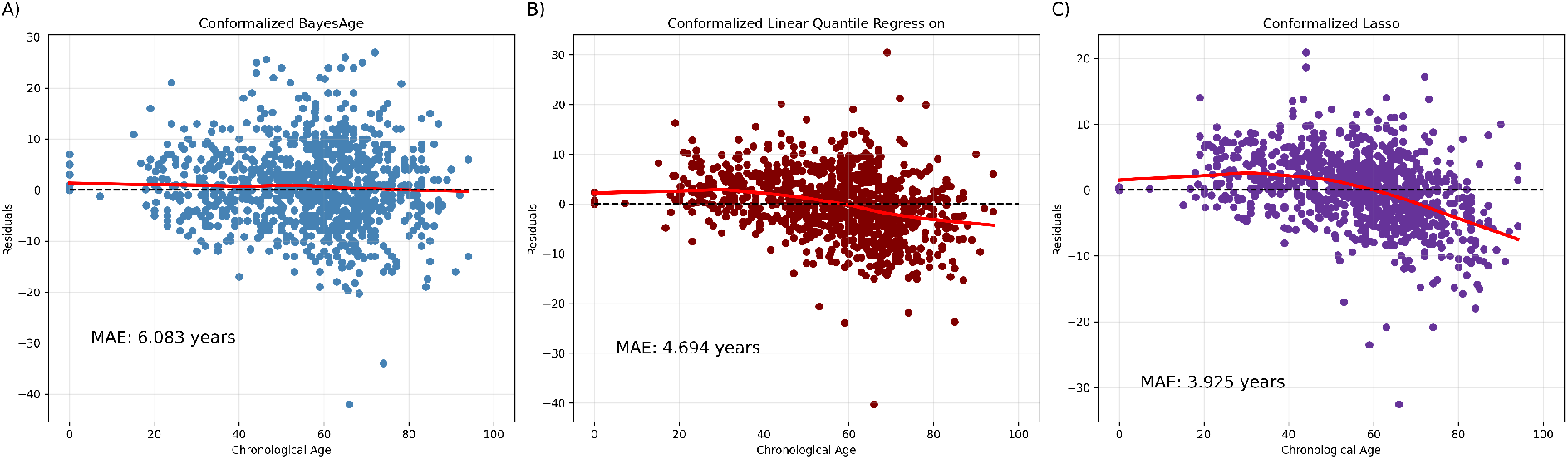
Signed residuals (age acceleration) versus chronological age for three predictors. **A)** Split conformalized BayesAge. **B)** Conformalized linear quantile regression. **C)** Split conformalized Lasso regression. Residuals are computed as predicted age minus chronological age. LOWESS smoothing summarizes age-dependent structure; the smoothing parameter is the software default.

## Discussion

We evaluated conformalized predictors of chronological age that attach prediction intervals to BayesAge point estimates. Under exchangeability between calibration and test examples, split conformal prediction provides finite-sample marginal coverage guarantees without requiring a parametric error model (15). This guarantee is marginal over draws of the calibration set and the test example; it does not imply conditional coverage given covariates such as age, tissue type, or study. In our setting, conformalization preserved BayesAge point-prediction accuracy while producing prediction intervals with near-nominal empirical coverage. Because BayesAge models nonlinear DNAm–age relationships using LOWESS reference curves and a count-based likelihood, this combination offers a practical route to uncertainty quantification for likelihood-based clocks that are not naturally expressed as standard regression models.

A key limitation of split conformal prediction is that it uses a single residual quantile per split and therefore yields constant-width intervals within that split prior to bounding. This can obscure age-dependent variability in prediction error that may arise from heteroskedastic DNAm dynamics and technical factors. To address this, we evaluated a locally scaled conformal variant that normalizes residuals by a locally estimated error scale and produces age-dependent interval widths (17, 18). In our data, locally scaled intervals tend to widen at older ages, consistent with increased absolute error in that regime. However, because the local scale must itself be estimated, finite-sample coverage can be less stable, particularly when calibration sets are small. Our small-calibration stress test illustrates this limitation. More stable local scale estimators, including spline-based approaches as in Lei et al. (17), may improve performance.

The simulation-based intervals used in the original BayesAge implementation substantially under-covered in this dataset. This behavior is consistent with a procedure that primarily propagates read-sampling variability conditional on observed DNAm fractions, while not directly accounting for model misspecification, cohort heterogeneity, or other sources of prediction error. In contrast, split conformal prediction calibrates intervals directly from held-out residuals and therefore adapts to the overall error distribution observed in the calibration data, without requiring an explicit generative model for that error.

We compared BayesAge-based intervals to higher-dimensional linear baselines wrapped in conformal prediction. In this study, conformalized Lasso regression produced narrower intervals and lower MAE than BayesAge, but required substantially more CpG sites. Conformalized linear quantile regression achieved intermediate widths and MAE at the cost of a larger feature set. BayesAge used far fewer CpG sites and, in this dataset and implementation, showed less age-dependent structure in residuals than the linear baselines. Because many applications focus on age acceleration, age-dependent residual patterns can complicate interpretation. We therefore view residual behavior as a complementary criterion alongside MAE and interval width. At the same time, residual structure is dataset- and implementation-dependent, and should be assessed whenever a clock is deployed in a new cohort.

This study also has limitations. The dataset was aggregated from multiple projects with diverse objectives, specimen types, and potential batch effects. Such heterogeneity can challenge exchangeability and can weaken formal marginal coverage guarantees when the deployment distribution differs from the calibration distribution; in practice, coverage should be assessed within relevant subgroups (for example, by tissue, study, or age range). We truncated reported intervals to the admissible age range [0, 100]. This truncation affects reported widths and can induce asymmetry near the boundaries, but it does not change whether the interval contains *Y* when *Y* ∈ {0, …, 100}. Several design choices, including the number of CpG sites and the smoothing parameter used for local scale estimation, were selected based on exploratory analyses in this cohort. A nested selection strategy or an external hold-out cohort would provide a cleaner separation between model selection and evaluation. Finally, because split conformal prediction provides marginal rather than conditional coverage, calibrated intervals may still under-cover within specific covariate strata, and this possibility should be checked in any deployment setting.

In summary, split conformal prediction provides a computationally simple, distribution-free approach to uncertainty quantification for DNAm-based age estimation, with finite-sample marginal coverage guarantees under exchangeability. Because it can wrap any point predictor, it can be applied to a broad class of epigenetic clocks, including likelihood-based models that capture nonlinear DNAm–age relationships. This can support applications in which quantified uncertainty is needed to interpret age estimates and age acceleration measures.

## Materials and Methods

### Data acquisition and processing

To evaluate uncertainty quantification for DNAm-based chronological age prediction, we analyzed targeted bisulfite sequencing (TBS-seq) data from 988 subjects. Samples were aggregated from multiple studies that used a common probe set enriched for CpG sites associated with age, cell-type composition, metabolic traits, and disease-related phenotypes (14, 19–24). Specimens included blood, buccal swab, and saliva. DNA extraction followed standard manufacturer protocols.

TBS-seq library preparation and capture were performed as described by Morselli et al. (25). Briefly, 250–500 ng of purified DNA per sample was used for library preparation. Fragmented DNA underwent end repair, dA-tailing, and adapter ligation using the NEBNext Ultra II library prep kit (New England Biolabs) with custom pre-methylated adapters (Integrated DNA Technologies). Libraries (16 per pool, each with unique adapters) were pooled and hybridized with biotinylated probes according to the manufacturer’s protocol. Captured DNA was bisulfite converted and PCR amplified using KAPA HiFi Uracil+ (Roche) under the following conditions: 2 minutes at 98^*°*^C; 14–16 cycles of 98^*°*^C for 20 seconds, 60^*°*^C for 30 seconds, and 72^*°*^C for 30 seconds; and 72^*°*^C for 5 minutes, followed by a hold at 4^*°*^C. Library quality control was performed using the High Sensitivity D1000 Assay on an Agilent 2200 TapeStation. Libraries were sequenced as paired-end 150 base reads on an Illumina NovaSeq 6000.

Raw reads were adapter- and quality-trimmed using `cutadapt`, aligned to the GRCh38 reference genome using `BSBolt Align`, and PCR duplicates were marked with `samtools markdup` . DNAm calls were generated using `BSBolt CallMethylation` and aggregated into a CpG-by-sample matrix using `BSBolt AggregateMatrix` . Matrices from individual studies were merged over common CpG sites, yielding a final matrix of 10,710 CpG sites across 988 samples. Software versions and key parameters used for read processing, modeling, and conformal calibration are reported as follows: `cutadapt v2.10`, `BSBolt v1.3.0, samtools v1.14`, `scikit-learn 1.8.0`, and `MAPIE 1.0.1` .

### BayesAge prediction model

We previously introduced BayesAge, a maximum likelihood estimation (MLE) method that predicts chronological age from DNAm counts and is based on the single-cell epigenetic clock scAge (26). BayesAge differs from high-dimensional linear regression approaches in two respects. First, it uses locally weighted scatterplot smoothing (LOWESS) to represent nonlinear relationships between DNAm fraction and age at each locus. Second, it models DNAm as count data using a binomial likelihood, which accommodates varying coverage across loci and can be less sensitive to missing counts (14).

BayesAge consists of a training step and a prediction step. During training, we filter CpG sites and retain those most strongly associated with age in the training data. Within each cross-validation split, we rank candidate sites using only the split’s training subset by the absolute Spearman rank correlation between DNAm fraction and chronological age, which is robust to monotone nonlinear trends. For that split, we then select the top *K* ranked CpG sites (here *K* = 16) and fit the BayesAge reference curves using only the training subset. The choice *K* = 16 was fixed for all experiments after exploratory analyses that evaluated MAE, empirical coverage, interval width, and computational cost (Fig. 10).

**Fig. 10.**
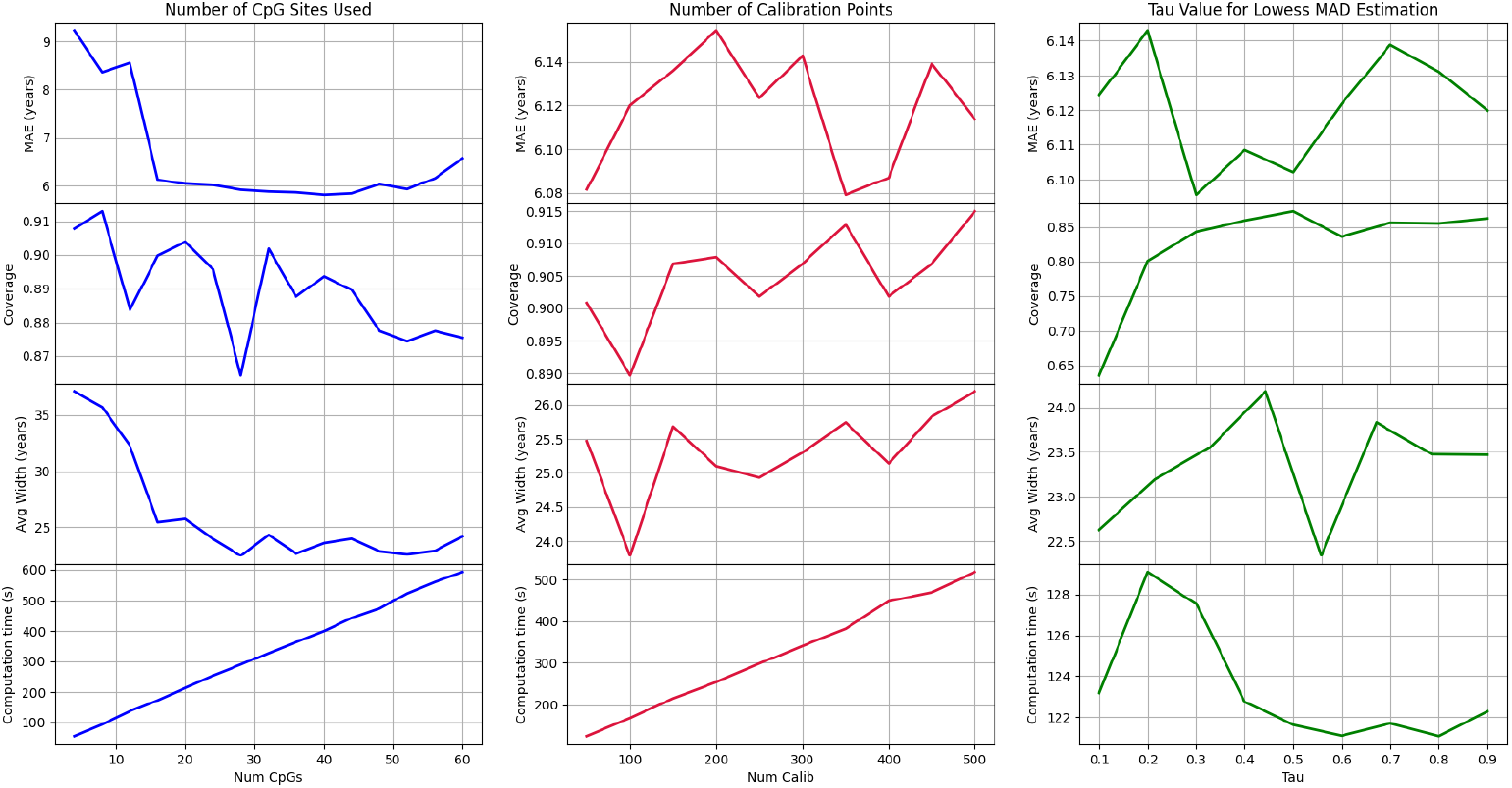
Hyperparameter exploration for conformalized BayesAge. We varied the number of CpG sites and the number of calibration samples in split conformal prediction, and we varied the LOWESS smoothing parameter *τ* used to estimate the local error scale for the locally scaled method.

For each selected CpG site *j* ∈ {1, …, 16}, we fit a LOWESS curve relating DNAm fraction to chronological age using the split’s training subset. LOWESS was fit with smoothing fraction 0.7. From each smoothed curve we extract the expected DNAm fraction at each integer age *a* ∈ *y*= {0, 1, …, 100} by evaluating the locally weighted regression at that point. These values define a deterministic reference matrix

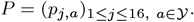

For prediction, let ℕ_0_ = {0, 1, 2, …}. Consider a sample with methylated read counts 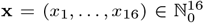and total read. depths 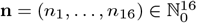 at the 16 loci. BayesAge evaluates, for each candidate age *a* ∈ *Y* a binomial likelihood in which loci are treated as conditionally independent given age:

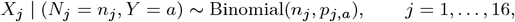

*X*_*j*_ | (*N*_*j*_ = *n*_*j*_, *Y* = *a*) ∼ Binomial(*n*_*j*_, *p*_*j,a*_), *j* = 1, …, 16, with conditional independence across *j* given (**N**, *Y*). The likelihood for candidate age *a* is

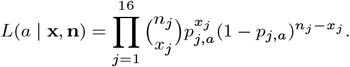

Equivalently, BayesAge maximizes the log-likelihood

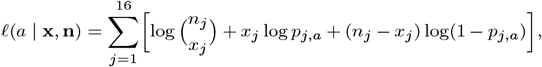

where boundary cases with *p*_*j,a*_ ∈{0, 1} are handled by clipping *p*_*j,a*_ to [*ϵ*, 1 − *ϵ*] with *ϵ* = 0.001 to avoid undefined logarithms. If a locus has *n*_*j*_ = 0 for a given sample, its contribution to *ℓ*(*a* **x, n**) is defined to be zero for all *a*, so that locus provides no information for the sample. The term 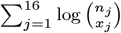 does not depend on a and can be omitted when computing the maximizer. The BayesAge point estimate is the maximizer over the discrete age grid,

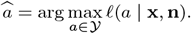

In practice, *ℓ* (*a* | **x, n**) is evaluated on the finite grid *Y*and the maximizing age is selected numerically. If ties occur, we break ties by selecting the smaller age. We use the binomial likelihood as a scoring function for point estimation on the discrete age grid; prediction intervals in this work are obtained by conformal calibration rather than by assuming a parametric distribution for the prediction error.

### Conformal prediction

Conformal prediction provides a general, model-agnostic approach to uncertainty quantification that can be wrapped around any point predictor (15). Under exchangeability of the labeled examples (a condition satisfied, for example, by independent and identically distributed samples), conformal predictors yield finite-sample marginal coverage guarantees. In contrast to methods that rely on large-sample approximations or parametric error models, conformal prediction calibrates prediction sets directly from held-out residuals and does not assume a specific distribution for the prediction error (27).

The central ingredient is a *nonconformity function*, a scalar score that measures how atypical a labeled example appears relative to a fitted predictor. A common choice is the absolute prediction error. In our setting, let ℕ_0_ = {0, 1, 2}, The feature (object) space is

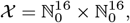

corresponding to methylated read counts and total read depths at the 16 selected CpG loci. BayesAge evaluates candidate ages on the integer grid

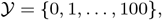

and throughout we report ages and residuals on this one-year grid. Let *Z*_*i*_ = *(X*_*i*_, *Y*_*i*_) ∈ *X* × *Y denote l*abeled examples, and assume {*Z*_*i*_}_*i*≥1_ is exchangeable.

### Full conformal prediction

Given training data (*x*_1_, *y*_1_), …, (*x*_*n*_ − _1_, *y*_*n*_ − _1_) and a test feature vector *x*_*n*_, full conformal prediction considers each candidate label *y* ∈ *Y by form*ing the augmented dataset

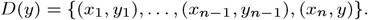

Let 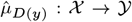 denote the BayesAge predictor fit on *D*(*y*). Using the absolute error as the nonconformity score, define

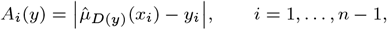

and

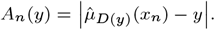

For a target coverage level *δ* ∈ (0, 1) and miscoverage *α* = 1 − *δ*, the full conformal prediction set is

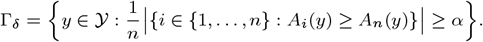

Under exchangeability, the standard conformal argument implies the marginal coverage guarantee

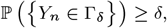

with the inequality accounting for discreteness and ties in the nonconformity scores. The set Γ_*δ*_ is a prediction set for the unknown chronological age *Y*_*n*_; when Γ_*δ*_ is an interval, its endpoints define a distribution-free (1 − *α*) prediction interval.

In practice, full conformal prediction is computationally burdensome in our setting because it requires refitting BayesAge for each candidate *y* ∈ *Y and for* each test sample. We therefore use split conformal prediction, which retains finite-sample marginal coverage under exchangeability while requiring only one fit of the predictor per data split.

### Split conformal prediction

Split conformal prediction is a computationally efficient alternative to full conformal prediction. The labeled data are split into a training set *D*_train_ and a calibration set *D*_cal_. The point predictor is fit once on *D*_train_, and prediction intervals are constructed using nonconformity scores computed on *D*_cal_.

Let 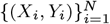be an exchangeable sequence with *X_i_* ∈ *X* and *Y_i_* ∈ *Y*. Partition the indices into disjoint sets *I_tr_* and *I_cal_*, and define

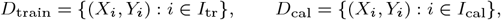

with *I*_tr_ ∩*I*_cal_ = ∅ and *m* = |*I*_cal_ |. Fit the BayesAge predictor 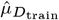 on *D*_train_, and compute calibration nonconformity scores using the absolute error

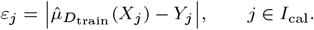

Let *ε*_(1)_ ≤ · · · ≤ *ε*_(*m*)_ denote the order statistics of {*ε*_*j*_ : *j* ∈ *I*_cal_}. For a target coverage level *δ* ∈ (0, 1), define

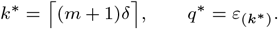

We use this standard non-randomized order-statistic definition; because the scores are discrete and can have ties, the resulting marginal coverage guarantee is generally conservative.

For a new feature vector *X*_new_ with point prediction 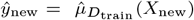, the split conformal prediction interval is

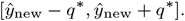

We interpret this as an interval on the real line, and we evaluate coverage by checking whether the true age *Y*_new_ ∈ Y {= 0, …, 100} lies within this interval.

Under exchangeability, the split conformal construction yields the finite-sample marginal coverage guarantee

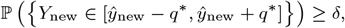

with the inequality accounting for discreteness and ties in the nonconformity scores.

We implemented split conformal prediction for BayesAge using code adapted from MLBoost (28). The full dataset contains 988 samples. We performed 12 splits. In each split, we fit a separate BayesAge predictor on a training subset, computed a conformal calibration quantile using a calibration subset, and evaluated coverage on a disjoint validation subset. We then report empirical coverage and interval-width summaries by pooling validation points across splits, so that each sample serves as a validation point exactly once. Under exchangeability within a split between calibration and test examples, split conformal prediction provides a finite-sample marginal coverage guarantee for the predictor defined by that split; the pooled empirical coverage summarizes performance across the collection of split-specific predictors.

For each split, we constructed the BayesAge reference matrix using the training set, computed calibration residuals *ε*_*j*_ on the calibration set, and selected *q*\* at rank *k*\* = ⌈ (*m* + 1) ⌉*δ* with *m* = 200. For each validation point with BayesAge prediction *ŷ*, we formed the unbounded split conformal interval [*ŷ* − *q*\*, *ŷ* + *q*\*]. For reporting, we bounded each interval to the admissible age range [0, 100]:

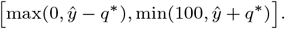

Because the label space is restricted to *Y {= 0*, …, 100}, this truncation affects reported widths and can induce asymmetry near the boundaries, but it does not change whether the interval contains the true age when the true age lies in *Y*.

### *Lo*cally scaled conformal prediction

Split conformal prediction uses a single calibration quantile per split and therefore produces constant-width intervals within that split prior to bounding. If prediction errors are heteroskedastic across age, constant-width intervals may not reflect meaningful changes in uncertainty. Heteroskedasticity is plausible in DNAm-based age prediction because individual CpG loci follow distinct DNAm trajectories and can exhibit age-dependent variability (13). DNAm is also influenced by environmental, lifestyle, and stochastic factors that can differ across age ranges and subpopulations (2). Motivated by these considerations, we evaluated a locally scaled conformal approach in which the interval half-width at a test point is proportional to a locally estimated error scale (17).

Conformal prediction permits general nonconformity scores. Let *D*_train_ and *D*_cal_ be a training/calibration split, and let 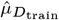denote the BayesAge predictor fit on *D*_train_. For calibration examples (*x*_*i*_, *y*_*i*_) ∈*D*_cal_, define point predictions 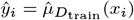 and a positive local scale estimate 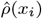. The locally scaled nonconformity score is

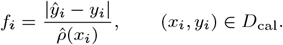

Let *f*_(1)_ ≤ · · · ≤ *f*_(*m*)_ denote the order statistics of the *m* = |*D*_cal_| calibration scores. For a target coverage level *δ* ∈ (0, 1), define

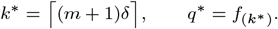

For a test object *x* with point prediction 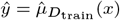, the locally scaled conformal interval is

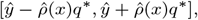

which we interpret as a real-valued interval and evaluate by checking inclusion of the integer-valued true age in *Y. The re*ported interval is obtained by bounding to the admissible range [0, 100] as in the split conformal procedure.

In our implementation, the local scale 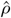is a function of predicted age. Specifically, we first compute the BayesAge point prediction *ŷ*_*i*_ for each calibration example and then estimate the conditional mean absolute deviation of the residual given predicted age via LOWESS smoothing. Concretely, we fit a LOWESS curve to the pairs (*ŷ*_*i*_, | *ŷ*_*i*_ − *y*_*i*_ |) on the calibration set and define 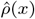 by evaluating the smoothed curve at the predicted age 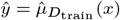. This choice allows nonlinear dependence of absolute error on predicted age. We tuned the LOWESS smoothing parameter *τ* over values in [0.1, 0.9] and used *τ* = 0.5 for the results reported here. We refer to *ŷ* as the predicted age.

Because 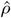 is estimated using calibration labels in this implementation, the standard split conformal coverage guarantee for the scaled scores does not strictly apply. A guarantee-preserving alternative is to estimate 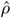 using training data only (or via cross-fitting) and then treat 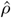 as fixed when computing the calibration scores *f*_*i*_.

#### Finite-sample coverage stress test

To probe finite-sample behavior in a regime where calibration is extremely limited, we performed a stress test with small splits. Specifically, we repeated 1000 random splits of 30 samples into 10 training, 10 calibration, and 10 validation examples, and evaluated both split conformal and locally scaled conformal prediction at target coverage *δ* = 0.9. For each split, empirical coverage was computed on the validation subset as the fraction of validation ages that fell within the reported prediction intervals. We report the running average of this coverage across splits to assess whether it approaches the nominal level.

### Comparison with alternative uncertainty quantification baselines

#### Monte Carlo simulation-based intervals

In the original BayesAge implementation, uncertainty was quantified by Monte Carlo simulation of DNAm counts. We applied the same procedure for comparison. For each individual and each CpG locus *j* ∈ {1, …, 16}, let *n*_*j*_ denote the observed total read depth and let *p*_*j*_ ∈ [0, 1] denote the observed DNAm fraction. Conditional on (*n*_*j*_, *p*_*j*_), we generated synthetic methylated counts according to

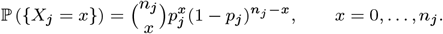

We repeated this sampling independently across the 16 loci and across 100 Monte Carlo draws to obtain 100 synthetic DNAm count vectors per individual.

For each synthetic vector, we predicted age using BayesAge and obtained, for each individual, a list of 100 simulated age predictions. We then formed a nominal 90% interval by taking the empirical 5th and 95th percentiles of this list. With 100 Monte Carlo draws, these percentiles are estimated with non-negligible Monte Carlo variability in the interval endpoints. Increasing the number of draws reduces this Monte Carlo error but does not address the main limitation of the procedure.

This procedure propagates read-sampling variability conditional on the observed DNAm fractions; it does not incorporate additional uncertainty sources such as cohort shift or model error. In addition, it uses a plug-in estimate of the methylation probability at each locus (*p*_*j*_ = *x*_*j*_ */n*_*j*_), which ignores uncertainty in the underlying methylation probability and any overdispersion beyond the binomial model (e.g., beta-binomial variability). Because BayesAge predictions are computed on a discrete age grid, the resulting Monte Carlo distribution of predicted ages can also be multi-peaked, in which case percentile intervals may be a poor summary of the prediction uncertainty.

#### Conformalized linear quantile regression baseline

We evaluated a conformalized linear quantile regression baseline using `MapieQuantileRegressor` with the default `QuantileRegressor` from `scikit-learn` (29, 30). We used the same calibration set size per split as in the conformalized BayesAge experiments, and we constructed prediction intervals at the same target coverage levels. CpG features for linear models were represented as DNAm fractions.

For feature selection, we varied the number of CpG sites from 8 to 256. Within each cross-validation split, we ranked CpG sites using only the split’s training subset by the absolute Spearman rank correlation between DNAm fraction and chronological age, and we retained the top *p* sites for each evaluated feature set size *p*. The same selected features for that split were then used to fit the quantile regression model on the training subset and to construct conformal intervals using the calibration subset. We observed diminishing improvements in MAE beyond 88 CpG sites while computation time increased with feature count. On this basis, we selected 88 CpG sites for the main conformalized linear quantile regression analysis.

#### Lasso regression baseline

Finally, we compared split conformalized BayesAge with split conformal prediction wrapped around Lasso regression as a high-dimensional linear baseline. Lasso is an *ℓ*_1_-penalized linear regression model that performs embedded feature selection and is conceptually similar to early linear epigenetic clocks.

We implemented Lasso using `scikit-learn` with a maximum of 10,000 iterations. CpG features were represented as DNAm fractions. To avoid notational conflict with the miscoverage level *α* = 1 − *δ*, we denote the Lasso penalty by *λ* (called `alpha` in `scikit-learn`). We tuned *λ* by cross-validation using the same splitting protocol as in the other baselines (with 88 validation samples for the first 11 splits and the final 20 samples in the last split). We evaluated *λ* ∈ [0.01, 1] and selected *λ* = 0.02, which minimized MAE under this protocol. At *λ* = 0.02, the fitted models used an average of 244.5 CpG sites with nonzero coefficients per split. We then wrapped these Lasso point predictors in the same split conformal procedure used for BayesAge to obtain prediction intervals and empirical coverage estimates.

## ACKNOWLEDGMENTS

We thank Michael Thompson for initial data curation, compilation, and cleaning.

## References

1. LD Moore, T Le, G Fan, DNA methylation and its basic function. Neuropsychopharmacology 38, 23–38 (2013).

2. G Hannum, et al., Genome-wide methylation profiles reveal quantitative views of human aging rates. Mol. Cell 49, 359–367 (2013).

3. L Vidal-Bralo, Y Lopez-Golan, A Gonzalez, Simplified assay for epigenetic age estimation in whole blood of adults. Front. Genet. 7, 126 (2016).

4. S Bocklandt, et al., Epigenetic predictor of age. PLOS ONE 6, e14821 (2011).

5. S Horvath, DNA methylation age of human tissues and cell types. Genome Biol. 14, 3156 (2013).

6. ME Levine, et al., An epigenetic biomarker of aging for lifespan and healthspan. Aging 10, 573–591 (2018).

7. RE Marioni, et al., DNA methylation age of blood predicts all-cause mortality in later life. Genome Biol. 16, 25 (2015).

8. Q Zhang, et al., Improved precision of epigenetic clock estimates across tissues and its implication for biological ageing. Genome Medicine 11, 54 (2019).

9. Q Lin, et al., DNA methylation levels at individual age-associated CpG sites can be indicative for life expectancy. Aging 8, 394–401 (2016).

10. D Kriukov, E Kuzmina, E Efimov, DV Dylov, EE Khrameeva, Epistemic uncertainty challenges aging clock reliability in predicting rejuvenation effects. Aging Cell 23, e14283 (2024).

11. RS Alisch, et al., Age-associated DNA methylation in pediatric populations. Genome Res. 22, 623–632 (2012).

12. S Snir, C Farrell, M Pellegrini, Human epigenetic ageing is logarithmic with time across the entire lifespan. Epigenetics 14, 912–926 (2019).

13. ND Johnson, et al., Non-linear patterns in age-related DNA methylation may reflect CD4+ t cell differentiation. Epigenetics 12, 492–503 (2017).

14. L Mboning, L Rubbi, M Thompson, LS Bouchard, M Pellegrini, BayesAge: A maximum likelihood algorithm to predict epigenetic age. Front. Bioinforma. 4, 1329144 (2024).

15. V Vovk, A Gammerman, G Shafer, Algorithmic learning in a random world. (Springer), Second edition edition, (2022).

16. Y Li, JM Goodrich, KE Peterson, PXK Song, L Luo, Uncertainty quantification in epigenetic clocks via conformalized quantile regression (2025).

17. J Lei, M G’Sell, A Rinaldo, RJ Tibshirani, L Wasserman, Distribution-free predictive inference for regression. J. Am. Stat. Assoc. 113, 1094–1111 (2018).

18. Y Romano, E Patterson, E. Candès, Conformalized quantile regression (2019).

19. J Chan, L Rubbi, M Pellegrini, DNA methylation entropy is a biomarker for aging. Aging 17, 685 (2025).

20. H Fu, et al., The response to influenza vaccination is associated with DNA methylation-driven regulation of t cell innate antiviral pathways. Clin. epigenetics 16, 114 (2024).

21. M Morselli, et al., DNA methylation profiles in pneumonia patients reflect changes in cell types and pneumonia severity. Epigenetics 17, 1646–1660 (2022).

22. W Guo, et al., Type-2 diabetes biomarker discovery and risk assessment through saliva DNA methylome. medRxiv pp. 2024–12 (2024).

23. G Protti, et al., The methylome of buccal epithelial cells is influenced by age, sex, and physiological properties. Physiol. Genomics 55, 618–633 (2023).

24. YL Chang, et al., Human DNA methylation signatures differentiate persistent from resolving mrsa bacteremia. Proc. Natl. Acad. Sci. 118, e2000663118 (2021).

25. M Morselli, et al., Targeted bisulfite sequencing for biomarker discovery. Methods 187, 13–27 (2021).

26. A Trapp, C Kerepesi, VN Gladyshev, Profiling epigenetic age in single cells. Nat. Aging 1, 1189–1201 (2021).

27. MLBoost, Uncertainty quantification (1): Enter conformal predictors (2024).

28. MT Rad, mtorabirad/MLBoost (2025) original-date: 2023-04-07T19:16:16Z.

29. T Cordier, et al., Flexible and Systematic Uncertainty Estimation with Conformal Prediction via the MAPIE library in Conformal and Probabilistic Prediction with Applications. (2023).

30. F Pedregosa, et al., Scikit-learn: Machine learning in Python. J. Mach. Learn. Res. 12, 2825–2830 (2011).

